# Self-assembled primary tumor clusters for precision cancer therapy

**DOI:** 10.1101/700724

**Authors:** Shenyi Yin, Ruibin Xi, Aiwen Wu, Jia-Fu Ji, Jianzhong Jeff Xi

## Abstract

Several patient-derived tumor models emerged recently as robust preclinical models. However, their potential to guide clinical therapy remained unclear. We report a novel model for personalized drug testing called patient-derived tumor-like cell clusters (PTC) that enabled us to accomplish personalized drug tests within 10-20 days. Mechanistically, PTCs result from the migration and aggregation of CD8+/CD44+ primary epithelial cells in a mixture of fibroblasts and macrophages, and form spheroid cultures. Phenotypic and genotypic profiling of PTCs showed a high degree of similarity with the original patient tumors, indicating that PTC structurally and functionally recapitulated original tumors. Over 200 PTC models have been established from at least six cancer types and diverse sampling approaches. Given its seamless integration with current clinical sampling approaches, the PTC platform will revolutionize chemotherapy programs for cancer patients.

## Introduction

Precision oncology seeks to identify effective therapeutic strategies for individual cancer patients. Omics technologies play a paramount role in precision medicine, including the identification of key modulators in signaling pathways and contributing to the understanding of cancers from statistical and macroscopic perspectives (Cristescu et al., 2015; Hoadley et al., 2018). However, the clinical application of omics technologies has been limited: a large proportion of cancer patients do not have any actionable mutations (Bailey et al., 2018), and among the patients with targetable genomic alterations, choosing the optimum treatment has been difficult due to the lack of effective theoretical models for the determination of dominant factors that can benefit patient clinical outcomes.

Increasing evidence has suggested that patient-derived tumor models can faithfully recapitulate human tumor biology and predict potential clinical responses. Patient-derived tumor xenografts (PDXs) can retain the idiosyncratic characteristics of different tumors from individual patient and measure drug efficacy more accurately than traditional methods (Byrne et al., 2017; Gao et al., 2015; Stewart et al., 2017). In addition, a group of 2D or 3D cell culture technologies have demonstrated their potential. Patient-derived tumor cells (PDC), mainly consisting of cancer stem cells, can retain the genomic and biological characteristics of and have shown concordance with the tumor of origin on clinical response. The 3D cultures and organoids cultured from healthy and tumor tissues from cancer patients have likewise made great progress (Pauli et al., 2017; Vlachogiannis et al., 2018). Patient-derived organoids (PDO) have been used to perform high-throughput drug screening and to model the treatment responses in metastatic gastrointestinal cancers (Crespo et al., 2017; van de Wetering et al., 2015a). Furthermore, organoids generated by human embryonic stem cells or induced pluripotent stem cells may circumvent the limited availability of high-quality human primary materials (Gjorevski et al., 2016; McCracken et al., 2014; Papapetrou, 2016; Spence et al., 2011).

Though promising, these models have several limitations. First, the central premise of personalized medicine is to facilitate patient-specific treatment decision-making within a clinically meaningful time window. Such a time window is generally less than 2-3 weeks for the determination of the best chemotherapy regimen for gastric and colorectal cancers pre- and post-operatively. However, both PDXs and PDO normally take more than a few months. Second, standardized cell culture conditions are needed to facilitate the dissemination and application of the model in clinical setting. PDO technologies demonstrated tissue-specific features (Fujii et al., 2016; Sachs et al., 2018; Sato et al., 2011), and huge efforts have to be made to develop, validate and standardize culture protocols for different resources. Third, the accuracy of the models on drug response assessment is of great importance. However, PDXs may undergo mouse-specific tumor evolution. PDC is subject to the drug-sensitivity variations and genetic and/or epigenetic alterations along the passages. PDO are exclusively composed of epithelial cells (Fujii et al., 2016; Sachs et al., 2018; van de Wetering et al., 2015a) and this intrinsic lack of stromal components may adversely influence the recapitulation of the drug-response pattern of the original tumor.

To address these limitations, we optimized the current culture medium and condition to maintain the integrity of the dissociated tumor primary cells including tumor epithelial cells, macrophages, fibroblasts, *etc*., and to generate as many tissue materials as possible within 2 weeks for drug sensitivity testing. Here, we established a new *in vitro* tumor model named patient-derived tumor-like cell clusters (PTC).

## Results

### Establishment of PTC *in Vitro* Tumor Models for Gastric and Colorectal Cancers

In 3D cultures and organoids, matrigel was added to model the *in vivo* tissue microenvironment. We hypothesized that cell cultures were enriched mainly through cell proliferation and differentiation in those 3D cultures and organoids, thus making it difficult to generate enough materials for personalized drug tests within one month. In order to maintain the integrity of tumor primary cell populations and accomplish the cell expansion and drug test within 2 weeks (**Figure 1A**), we removed matrigel and emphasized a strategy of tumor cells self-assembly and proliferation. We tested a variety of culture media and polymers with good biocompatibility for the establishment gastric and colorectal cancer PTC. The substrate hydrophobicity appeared to have a substantial effect on the growth of PTC (**Figure S1A**). PTC demonstrated distinct phenotypes from PDOs (**Figure S1B**). For case CS-T-858, PTC presented compact clusters devoid of a lumen, whereas PDOs showed thin-walled cystic structures. In general, PTC had more but smaller clusters than PDOs.

**Figure 1.**
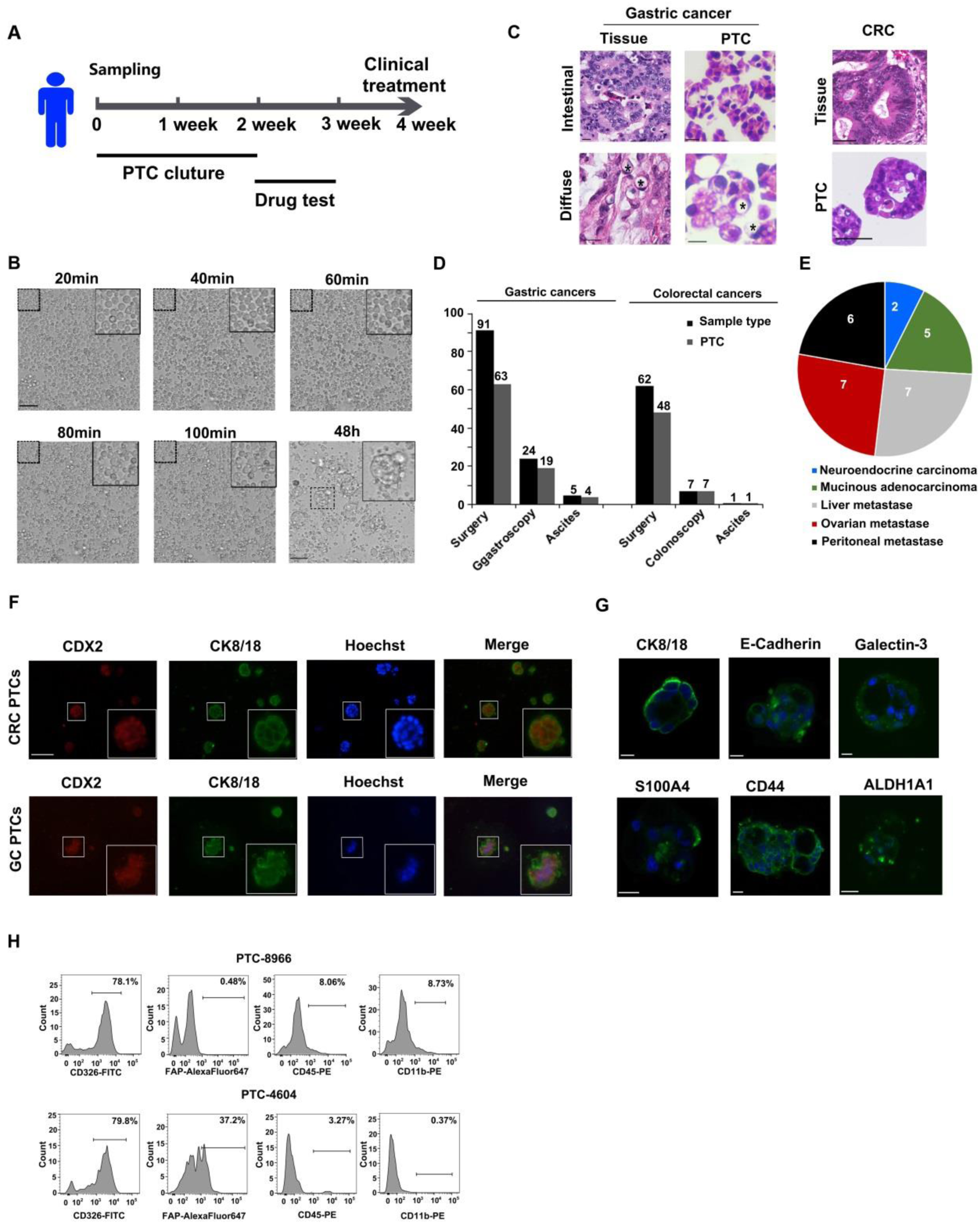
Generation of patient-derived tumor-like cell clusters (PTC) within 2 weeks. (A) An overview of the timeline of the PTC culture and personalized drug tests. (B) A time-lapse imaging sequence of the primary cell clustering in the first 100 minutes and 48 hours. Scale bars: 50 μm. (C) Hematoxylin and eosin (H&E) staining comparing the PTC to their matching patient biopsies. * indicates signet cells. Scale bar: 20 µm. (D) The numbers of different gastric or colorectal cancer samples collected and used to successfully generate PTC. The samples included surgically resected tissues, endoscopically obtained tissues and ascites tissues. (E) Distribution of gastric or colorectal cancers-derived PTC based on cancer subtypes, including neuroendocrine carcinoma, mucinous adenocarcinoma, or metastasis. (F) Immunofluorescent co-staining of the colorectal cancer (CRC) or gastric cancer (GC) marker CDX2 and the epithelial markers CK8/18. Nuclear counterstaining (blue, Hoechst). Scale bar: 100 µm. (G) Immunofluorescent staining of day 5 PTC for the epithelial markers CK8/CK18, the cell-cell junction marker E-cadherin (ECAD), the macrophage marker galectin-3, the fibroblast marker S100A4, the stem cell markers CD44 and ALDH. Nuclear counterstaining (blue, Hoechst). Scale bars: 20 μm. (H) Quantitative analysis of the proportion of epithelial cells (CD326+), fibroblast cells (FAP+) or macrophage cells (CD45+ or CD11b+) in two PTC samples (patient 8966 and 4604) by flow cytometry. See also **Figure S1** and **Movie S1.**

We dynamically studied the generation of PTC by a high-content imaging system (MD) and found that PTCs mainly resulted from the migration and self-assembly of dissociated primary cells into hundreds of clusters (**Figure 1B**). This finding is consistent with the substrate hydrophobicity mentioned above, as a hydrophilic surface may inhibit the assembly of PTC due to the adherent interaction between the cells and substrates. Furthermore, the PTC has demonstrated high morphological similarities with the gastrointestinal cancer biopsies (**Figure 1C**). For example, GS-2281 PTC, which was derived from diffuse-type gastric carcinoma, reconstituted the histologic appearance of signet ring cells as the original parental tumor.

A total of 190 fresh gastric or colorectal tumor samples including 153 surgically-resected tumor tissues, 31 endoscopic biopsies and 6 ascites samples were collected for PTC establishment. The overall success rate of PTC establishment was 74.7% (142/190). Although biopsy samples obtained through endoscopy had a smaller amount of tissue, the corresponding PTC had a higher success rate than that of those generated from surgically resected samples (83.9% vs 72.5%) (**Figure 1D**). Furthermore, the PTC can be easily adapted to study cancer subtypes for which it has historically been impossible or difficult to establish organoids (Fujii et al., 2016), such as poorly or moderately differentiated adenocarcinoma, neuroendocrine carcinoma, mucinous adenocarcinoma, or metastatic samples (**Figure 1E and S1C**).

### Characterization of PTC Cellular Components

We next investigated the cellular components of PTC prior to the genomic characterization. The immunofluorescence data revealed that the majority of the clustering cells were CK8/18+ epithelial cells (**Figure 1F, 1G**). Besides epithelial cell markers, PTC exhibited positive staining of tissue-specific markers. CRC PTC showed the expression of MUC2 and CDX2 (Crespo et al., 2017; Fujii et al., 2016; Kodack et al., 2017; van de Wetering et al., 2015b), while the PTC derived from gastric cancer samples showed the expression of CDX2, PDX1 and SOX2 (Kodack et al., 2017; McCracken et al., 2014). Similar to PDC and PDO, most PTC demonstrated the properties of cancer stem cells and high proliferation capability, as evidenced by CD133+/ALDH1A1+/Ki-67+ (**Figure 1G** and **S1D**).

Strikingly, PTC contained macrophages or fibroblast cells (**Figure 1G, S1E**). Real-time PCR assays showed that a large proportion of CD68+/CD163+ macrophages or S100A4+/α-SMA+ fibroblasts were detected in 43 individual clusters. Furthermore, we used flow cytometry assays to quantitatively analyze the proportion of epithelial, macrophages or fibroblast cells in seven PTC. We found that the proportions of non-epithelial cells varied significantly among patients, with fibroblast cells ranging from 0% to 50.7% and macrophage cells from 0% to 9% (**Figure 1H, S1F**).

### Genomic Consistency of PTC with the Original Tumors

Genomic DNA was isolated from the PTC cells, tumor samples and matched normal samples for target sequencing and low-pass whole genome sequencing (WGS). Somatic mutations and copy number variations (CNV) were called based on target sequencing and the low-pass WGS, respectively. Both of them were concordant between the PTC cells and the tumor samples. The proportions of common somatic mutations ranged from 0.57 to 1, with a mean of 0.86 and a standard deviation (s.d.) of 0.13 (**Figure 2A** and **S2**). Correlations of the CNVs between the PTC cells and tumor samples ranged from 0.52 to 0.96, with a mean of 0.78 and a s.d. of 0.13. Many samples had nearly identical CNV patterns between the PTC cells and tumor samples (**Figure 2B**). The observed minor differences in genomic alterations between the PTC cells and tumor samples could be *bona fide* genomic differences or artifacts caused by insufficient detection power. For example, common low frequency subclonal mutations usually have a lower detection power and are more likely to be misclassified as private mutations in PTC cells or tumor samples. In fact, when we increased the variant allele frequency to at least 20%, the mean proportion of common somatic mutations increased to 0.93 with a s.d. of 0.17.

**Figure 2.**
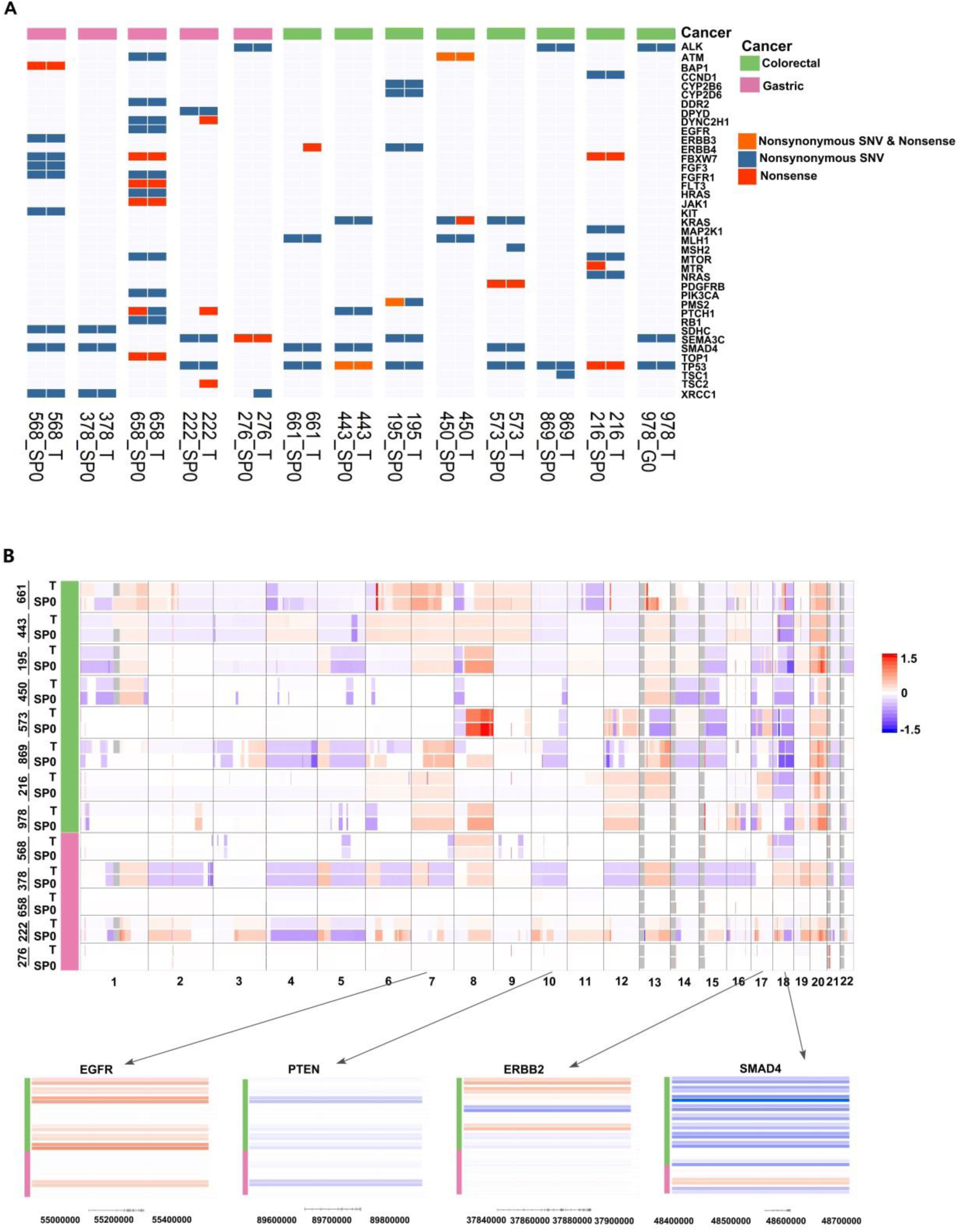
Genomic comparison between tumor biopsies and PTC. (A) Overview of the identified somatic mutations. Nonsense mutations included all nonsense SNVs, SNVs at splicing-sites and frameshift indels. Nonsynonymous mutations include nonsynonymous SNVs and in-frame indels. Nonsynonymous & Nonsense means the gene had both nonsense and nonsynonymous mutations. T stands for tissue; Sp0 stands for the spheres or clusters of PTC in different wells. (B) Overview of CNVs. The rows represented different samples and the columns were genomic positions from chromosome 1 to 22. Colors in the plot corresponded to the estimated log2 copy ratio of the genomic regions. The four bottom panels were local views of four cancer genes frequently altered in colorectal and gastric cancers. See also **Figure S2**.

The discovered genomic alternations largely agreed with previously reported mutations that were typical for colorectal and gastric cancers. Among the genes covered in our gene panel, *TP53, KRAS, PIK3CA, SMAD4* and *NRAS* were the top 5 most frequently mutated driver genes in colorectal cancer (Cancer Genome Atlas, 2012), and *TP53, SMAD4* and *PIK3CA* were the top 3 most frequently mutated genes in gastric cancer (Cancer Genome Atlas Research, 2014). Of the eight colorectal cancer patients in this study, seven had *TP53* non-synonymous mutations, three had *KRAS* mutations, zero had *PIK3CA* mutations, three had *SMAD4* mutations and one had a *NRAS* mutation. In these five genes, the PTC cells and the corresponding tumor tissues had the same exactly same mutations. For the five gastric cancer patients, we observed one patient, two patients and one patient with mutations in *TP53, SMAD4* and *PIK3CA*, respectively. The PTC cells and tumor samples from these patients also shared the same mutations in the respective genes. The typical arm-level CNVs in colorectal and gastric cancer were observed (Cancer Genome Atlas, 2012; Cancer Genome Atlas Research, 2014). For example, many samples had copy number gains of chromosomes 1q, 7, 8q, 13q, and 20, as well as copy number losses of 1p, 5q, 8p, 14q, 15q, 17p and 18. Recurrent focal amplifications and deletions were also observed in many patients, including amplifications of *ERBB2* and *EGFR*, and deletions of *PTEN* and *SMAD4* (**Figure 2B**). In summary, these genomic analyses revealed that the PTC faithfully maintained the genomic features of the primary tumors.

### PTC as a Tool for Personalized Drug Testing

To identify the optimum therapeutic option for individual patients, we set up a personalized drug testing system. The PTC were separated into a multi-well chip and the drugs to be evaluated were added into different individual wells. The cell clusters were photographed at day 0 and day 7. Only clusters with diameters of more than 40 micrometers were selected to calculate the cluster areas. The efficacy of a drug A was calculated by the following formula,

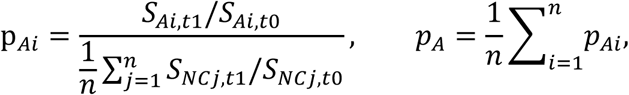

where S is the sum of cluster areas in a well, subscript NC is the negative control and used for normalization, n is the number of replication for the treatment and control, t0 and t1 are the time points when the areas are measured. On the basis of the RECIST criteria of solid tumor response used in clinical setting (Eisenhauer et al., 2009), the efficacy was divided into the following five categories:no effect, p_A_ ≥ 0.7; weak effect 0.7 > p_A_ ≥ 0.6; modest effect, 0.6 > p_A_ ≥ 0.4; significant effect, 0.4 > p_A_ ≥ 0.2; strong effect, p_A_ < 0.2 (**Figure S3A**).

We next evaluated whether PTC could be applied as an informative tool for precision oncology using the following two dimensions: the consistency among PTC in different wells and that between PTC’ prediction results on drug response and the patients’ actual responses in the clinical setting. The former consistency was addressed using two approaches: the drug-efficacy consistency and the genomic consistency.

For the drug-efficacy consistency, we firstly investigated the effect of the cluster numbers on the drug response consistency (**Figure 3A**). We then performed drug efficacy tests for 28 samples using 5-20 drugs and repeated the experiment three times for each drug-sample pair. As a result, we obtained 339 drug profiles in total. Drug efficacy consistency requires that different experiments for the same drug-sample pair render close and highly-correlated results. We calculated the CV of the three experiments for each drug-sample pair and found that most (91.74%) of these CVs were less than 0.5 (**Figure 3B**), indicating that the result of three experiments for most drug-sample pairs were very close to one another. In addition, we randomly selected two experiments for each drug-sample pair and calculated their Pearson’s correlation coefficient. This process was repeated for 1,000 times. We found that the correlations were consistently larger than 0.8, with a mean of 0.84 and a s.d. of 0.036 (**Figure 3C** and **S3B**), which indicated that results rendered by different experiments for the same drug-sample pair were highly consistent.

**Figure 3.**
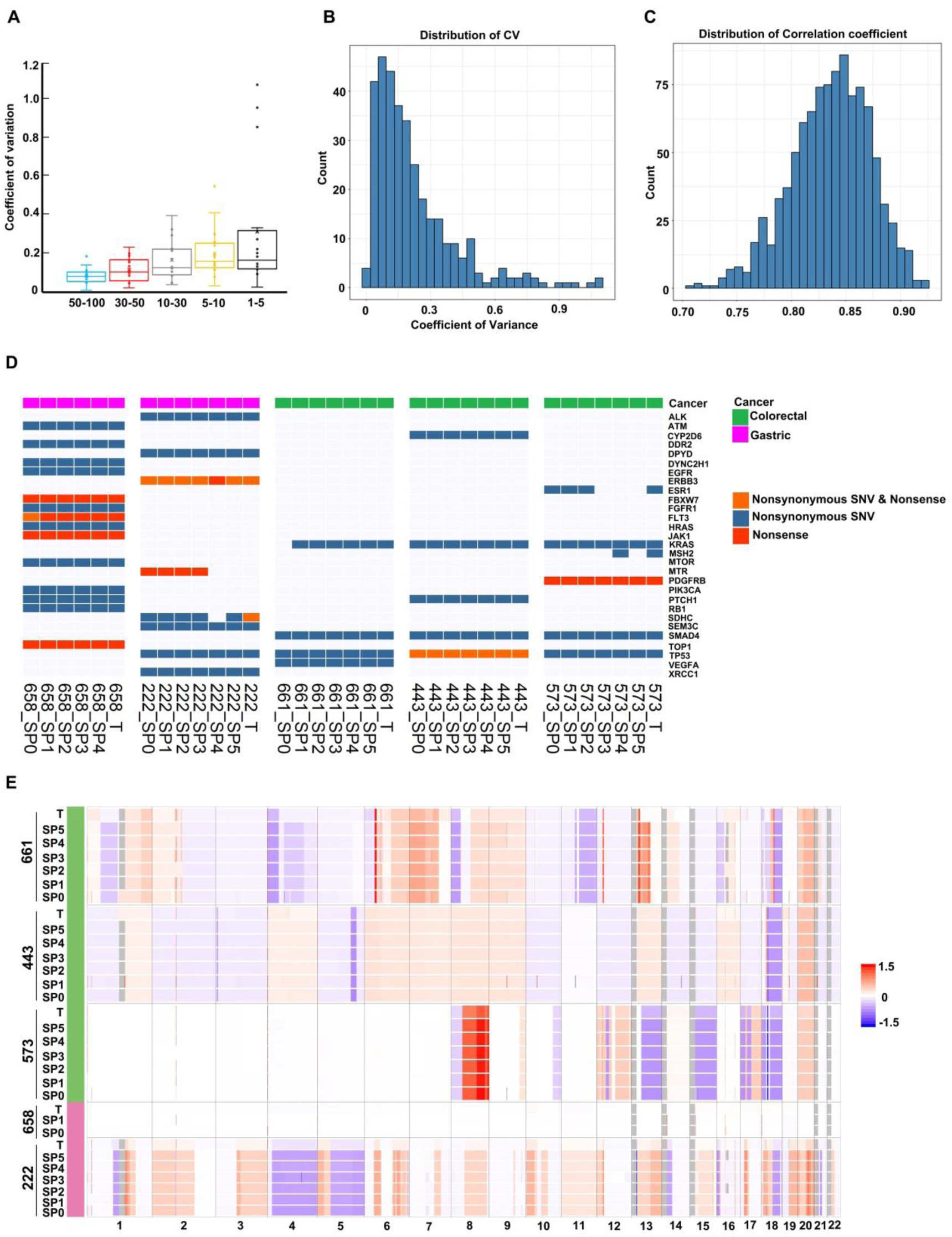
The establishment of the PTC system for personalized drug testing. (A) The effect of the cluster numbers on the drug response consistency. The y-axis is the coefficient of variation (CV) and x-axis is the number of clusters used. (B) The distribution of CVs in the drug-sample pairs in the gastrointestinal group. n = 339. (C) The distribution of the Pearson correlation coefficients in a 1000-fold random sampling strategy for two results in three parallel drug sensitivity tests. n = 339. (D, E) The somatic mutations and CNVs in tumors and in replicated PTC. T: tumor; SP0-SP5: the PTC in different wells. See also **Figure S3**.

We then investigated the genomic consistency in different wells by comparing the genomic and transcriptomic profiles of PTC. The somatic mutations, CNVs, and transcriptome profiles showed high consistency, indicating the similarity of the proportions and compositions of the PTC among wells (**Figure 3D, 3E and S3C**). With the exception of sample 222, pairwise overlap analysis showed that the proportions of common somatic mutations between different wells were always greater than 0.8 (**Figure S3D**). The CNV correlations between the cells had a mean of 0.91 and a s.d. of 0.09 (**Figure S3E**). The gene expression levels between different wells were likewise highly correlated, with many pairs of wells having correlations close to 1 (**Figure S3F**).

## Discussion

Time and accuracy are two paramount factors to develop any i*n vitro* tumor models that can provide valuable reference for clinician to make prescription. Here, we reported a novel personalized drug testing technology called PTC, and proved the effectiveness of the PTC experimental paradigm for individualized drug tests within 10-20 days, even starting with trace biopsies. In comparison with recently developed 3D organoids technologies (Drost and Clevers, 2018), the PTC technology demonstrates unique characteristics in manufacturing methods, organizing mechanism and composition, more importantly in time and accuracy. PTC established clear culture conditions, drug efficacy concentrations, and efficiency cut-off for accurate clinical prediction (**Figure S4**).

One counterintuitive observation is that organoids from normal epithelium often grow faster than organoids derived from advanced tumors (Drost et al., 2015; Verissimo et al., 2016). Various chemicals have to be added or removed to inhibit the growth of healthy cells or to promote culturing of different subtypes of tumor epithelial cells. Thus, extra time and efforts are needed to validate the accuracies of those organoids prior to their clinical applications. More severely, organoid cultures generally require a few months to generate sufficient materials for drug efficacy testing.

In this work, we tried to maintain the histological and functional integrity of tumor tissues by taking advantage of tumor’s self-organizing capacity. We found that the PTC resulted from the migration and self-assembling of dissociated tumor primary cells. We confirmed that, in addition to CK8/18+ epithelial cells, the PTC contained stromal cells such as microphage and fibroblast cells. These stromal cells played an indispensable role in PTC assembling and drug-response recapitulation.

Unlike the organoid cultures, in which different tumor subtypes require specific compositions of culture media (Drost and Clevers, 2018; Fujii et al., 2016), the PTC could be easily adapted to characterize drug sensitivity of other cancers besides gastric and colorectal cancers, and even a diversity of cancer subtypes that were impossible or difficult to generate corresponding organoids, such as poorly or moderately differentiated adenocarcinoma, neuroendocrine carcinoma, mucinous adenocarcinoma, or metastatic samples. More importantly, the PTC can forecast patients’ clinical responses to targeted agents or chemotherapies within 10-20 days. Although both PDX and PDO technologies retrospectively demonstrated the success in forecasting the patients’ drug response, whether they are applicable to the clinic in a timely manner remains to be determined (Gao et al., 2015; Vlachogiannis et al., 2018). Since the entire evaluation process of PTC can be seamlessly integrated into neoadjuvant, chemotherapy or target therapy programs, we strongly believe that PTC would revolutionize precision oncology and benefit millions of cancer patients every year.

## Reference

Bailey, M. H., Tokheim, C., Porta-Pardo, E., Sengupta, S., Bertrand, D., Weerasinghe, A., Colaprico, A., Wendl, M. C., Kim, J., Reardon, B., et al. (2018). Comprehensive Characterization of Cancer Driver Genes and Mutations. Cell 173, 371–385 e318.

Byrne, A. T., Alferez, D. G., Amant, F., Annibali, D., Arribas, J., Biankin, A. V., Bruna, A., Budinska, E., Caldas, C., Chang, D. K., et al. (2017). Interrogating open issues in cancer precision medicine with patient-derived xenografts. Nat Rev Cancer 17, 254–268.

Cancer Genome Atlas, N. (2012). Comprehensive molecular characterization of human colon and rectal cancer. Nature 487, 330–337.

Cancer Genome Atlas Research, N. (2014). Comprehensive molecular characterization of gastric adenocarcinoma. Nature 513, 202–209.

Crespo, M., Vilar, E., Tsai, S. Y., Chang, K., Amin, S., Srinivasan, T., Zhang, T., Pipalia, N. H., Chen, H. J., Witherspoon, M., et al. (2017). Colonic organoids derived from human induced pluripotent stem cells for modeling colorectal cancer and drug testing. Nat Med 23, 878–884.

Cristescu, R., Lee, J., Nebozhyn, M., Kim, K. M., Ting, J. C., Wong, S. S., Liu, J., Yue, Y. G., Wang, J., Yu, K., et al. (2015). Molecular analysis of gastric cancer identifies subtypes associated with distinct clinical outcomes. Nat Med 21, 449–456.

Drost, J., and Clevers, H. (2018). Organoids in cancer research. Nat Rev Cancer.

Drost, J., van Jaarsveld, R. H., Ponsioen, B., Zimberlin, C., van Boxtel, R., Buijs, A., Sachs, N., Overmeer, R. M., Offerhaus, G. J., Begthel, H., et al. (2015). Sequential cancer mutations in cultured human intestinal stem cells. Nature 521, 43–47.

Eisenhauer, E. A., Therasse, P., Bogaerts, J., Schwartz, L. H., Sargent, D., Ford, R., Dancey, J., Arbuck, S., Gwyther, S., Mooney, M., et al. (2009). New response evaluation criteria in solid tumours: revised RECIST guideline (version 1.1). Eur J Cancer 45, 228–247.

Fujii, M., Shimokawa, M., Date, S., Takano, A., Matano, M., Nanki, K., Ohta, Y., Toshimitsu, K., Nakazato, Y., Kawasaki, K., et al. (2016). A Colorectal Tumor Organoid Library Demonstrates Progressive Loss of Niche Factor Requirements during Tumorigenesis. Cell Stem Cell 18, 827–838.

Gao, H., Korn, J. M., Ferretti, S., Monahan, J. E., Wang, Y., Singh, M., Zhang, C., Schnell, C., Yang, G., Zhang, Y., et al. (2015). High-throughput screening using patient-derived tumor xenografts to predict clinical trial drug response. Nat Med 21, 1318–1325.

Gjorevski, N., Sachs, N., Manfrin, A., Giger, S., Bragina, M. E., Ordonez-Moran, P., Clevers, H., and Lutolf, M. P. (2016). Designer matrices for intestinal stem cell and organoid culture. Nature 539, 560–564.

Hoadley, K. A., Yau, C., Hinoue, T., Wolf, D. M., Lazar, A. J., Drill, E., Shen, R., Taylor, A. M., Cherniack, A. D., Thorsson, V., et al. (2018). Cell-of-Origin Patterns Dominate the Molecular Classification of 10,000 Tumors from 33 Types of Cancer. Cell 173, 291–304 e296.

Kodack, D. P., Farago, A. F., Dastur, A., Held, M. A., Dardaei, L., Friboulet, L., von Flotow, F., Damon, L. J., Lee, D., Parks, M., et al. (2017). Primary Patient-Derived Cancer Cells and Their Potential for Personalized Cancer Patient Care. Cell Rep 21, 3298–3309.

McCracken, K. W., Cata, E. M., Crawford, C. M., Sinagoga, K. L., Schumacher, M., Rockich, B. E., Tsai, Y. H., Mayhew, C. N., Spence, J. R., Zavros, Y., and Wells, J. M. (2014). Modelling human development and disease in pluripotent stem-cell-derived gastric organoids. Nature 516, 400–404.

Papapetrou, E. P. (2016). Patient-derived induced pluripotent stem cells in cancer research and precision oncology. Nat Med 22, 1392–1401.

Pauli, C., Hopkins, B. D., Prandi, D., Shaw, R., Fedrizzi, T., Sboner, A., Sailer, V., Augello, M., Puca, L., Rosati, R., et al. (2017). Personalized In Vitro and In Vivo Cancer Models to Guide Precision Medicine. Cancer Discov 7, 462–477.

Sachs, N., de Ligt, J., Kopper, O., Gogola, E., Bounova, G., Weeber, F., Balgobind, A. V., Wind, K., Gracanin, A., Begthel, H., et al. (2018). A Living Biobank of Breast Cancer Organoids Captures Disease Heterogeneity. Cell 172, 373–386 e310.

Sato, T., Stange, D. E., Ferrante, M., Vries, R. G., Van Es, J. H., Van den Brink, S., Van Houdt, W. J., Pronk, A., Van Gorp, J., Siersema, P. D., and Clevers, H. (2011). Long-term expansion of epithelial organoids from human colon, adenoma, adenocarcinoma, and Barrett’s epithelium. Gastroenterology 141, 1762–1772.

Spence, J. R., Mayhew, C. N., Rankin, S. A., Kuhar, M. F., Vallance, J. E., Tolle, K., Hoskins, E. E., Kalinichenko, V. V., Wells, S. I., Zorn, A. M., et al. (2011). Directed differentiation of human pluripotent stem cells into intestinal tissue in vitro. Nature 470, 105–109.

Stewart, E., Federico, S. M., Chen, X., Shelat, A. A., Bradley, C., Gordon, B., Karlstrom, A., Twarog, N. R., Clay, M. R., Bahrami, A., et al. (2017). Orthotopic patient-derived xenografts of paediatric solid tumours. Nature 549, 96–100.

van de Wetering, M., Francies, H. E., Francis, J. M., Bounova, G., Iorio, F., Pronk, A., van Houdt, W., van Gorp, J., Taylor-Weiner, A., Kester, L., et al. (2015a). Prospective derivation of a living organoid biobank of colorectal cancer patients. Cell 161, 933–945.

van de Wetering, M., Francies, H. E., Francis, J. M., Bounova, G., Iorio, F., Pronk, A., van Houdt, W., van Gorp, J., Taylor-Weiner, A., Kester, L., et al. (2015b). Prospective derivation of a living organoid biobank of colorectal cancer patients. Cell 161, 933–945.

Verissimo, C. S., Overmeer, R. M., Ponsioen, B., Drost, J., Mertens, S., Verlaan-Klink, I., Gerwen, B. V., van der Ven, M., Wetering, M. V., Egan, D. A., et al. (2016). Targeting mutant RAS in patient-derived colorectal cancer organoids by combinatorial drug screening. Elife 5.

Vlachogiannis, G., Hedayat, S., Vatsiou, A., Jamin, Y., Fernández-Mateos, J., Khan, K., Lampis, A., Eason, K., Huntingford, I., Burke, R., et al. (2018). Patient-derived organoids model treatment response of metastatic gastrointestinal cancers. Science 359, 920–926.

