## Supplementary data for "Self-assembled primary tumor clusters for precision cancer therapy"

**Abbreviation of Chemicals**

- **A**fatinib (Afa); Alectinib (Ale); Apatinib (Apa);
- Carboplatin (Car or Ca); Cetuximab (Cet); Cisplatin (Cis or Ci); Crizotinib (Cri);
- Docetaxel (Doc or D); Dovitinib (Dov)
- Epirubicin (Epi); Etoposide (Eto);
- Fluorouracil or Fluoropyrimidine (F); Folinate Calcium (Fol);
- Gefitinib (Gef); Gemcitabine (Gem or G);
- Icotinib (Ico); Irinotecan (Iri);
- Mitomycin (Mit or M)
- Olaparib (Ola); Osimertinib (Osi); Oxaliplatin (Oxa);
- Paclitaxel (Pa); Pemetrexed (Pem or Pe); Pertuzumab (Per); Pirarubicin (Pir); Ponatinib (Pon)
- Raltitrexed (Ral); Regorafenib (Reg)
- Sorafenib (Sor)
- Trametinib (Tra); Trastuzumab (Tra);
- Vemurafenib (Vem); Vinorelbine (Vin or V)

#### Developing PTC as a Tool for Personalized Drug Testing

To identify the optimum therapeutic option for individual patients, we designed a personalized drug testing system based on PTCs: we separate thousands of PTCs into a multi-well chip and evaluate drugs in different wells. To inform treatment design, we evaluated drug effect, determined efficacy cut-off, and found the drug efficacy concentration (**Figure S4**).

First, we evaluated drug effect by measuring the area of all PTC clusters in a well. The cell clusters were photographed at day 0 and day 7 in a well. Only clusters with diameters of more than 40 micrometers were selected to calculate the cluster areas. The efficacy of a drug A was calculated by the following formula,

$$\mathrm{p}_{Ai} = \frac{S_{Ai,t1}/S_{Ai,t0}}{\frac{1}{n}\sum_{j=1}^{n} S_{NCj,t1}/S_{NCj,t0}}, \qquad p_A = \frac{1}{n}\sum_{i=1}^{n} p_{Ai},$$

where S is the sum of cluster areas in a well, subscript NC is the negative control and used for normalization, n is the number of replications for the treatment and control, and t0 and t1 are the time points when the areas are measured.

Second, we determined the clinical efficacy of a drug A among patients according to the RECIST criteria (Eisenhauer et al., 2009). For the purpose of analysis, we collapsed the RECIST criteria into two groups using 0.7 as cut-off, and defined complete response (CR) or partial response (PR) as effective, and stable disease (SD) or progressive disease (PD) as otherwise. Accordingly, the efficacy was divided into the following two categories: the drug was not effective if $\mathrm{p_A} \geq 0.7$; effective if $\mathrm{p_A} < 0.7$ (**Figure S4**).

Third, we determined the concentration of the drug added to the wells, i.e. efficacy concentration (Ec), according to its clinical efficacy. If the PTC model would accurately predict the individual's clinical response, the efficacy rate of any drug would be consistent with the overall response rate of this drug among patients. The concentration-response profile of a given drug A was generated using 20-30 patients' PTC samples. We set the Ec as the concentration such that the effective rate in PTC drug test was closest to the overall response rate of drug A previously reported in relevant clinical trials (**Figure S4**). For example, a group of clinical trials showed that overall response rate was about 11%- 20% for 5-fluorouracil in advanced GC or CRC (Cunningham et al., 1996; Liu et al., 2014; Ohtsu et al., 2003). Accordingly, 5-Fluorouracil Ec was set to 2 μmol/L, at which the efficacy rate was 18.1% (6/33).

### Supplementary Figure legends

**Figure S1**

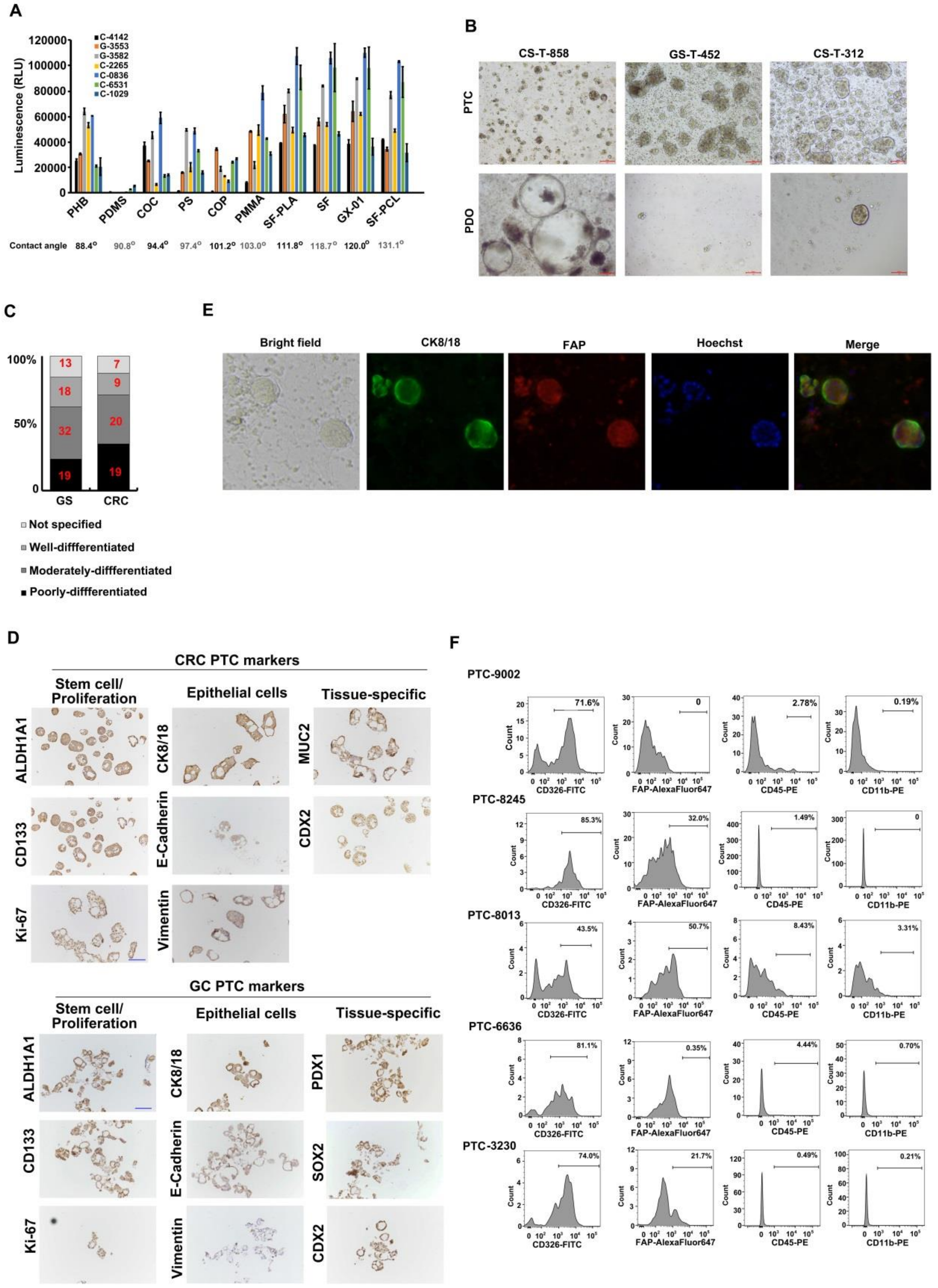


**Figure S1**. **Culture and Characterization of PTC, Related to Figure 1.**

(A) The effect of culture substrates on the PTC growth. The hydrophobic property of each substrate was measured as a contact angle. PHB: Polyhydroxybutyrate; PDMS: polydimethylsiloxane; COC: Cyclic olefin copolymer; PS: Polystyrene; COP: Cyclo Olefin Polymer; PMMA: Poly(methyl methacrylate); SF-PLA: silk fibroin-poly (L-Lactic acid); SF: silk fibroin (Shi et al., 2017); GX-01(GeneX); SF-PCL: silk fibroin polycaprolactone (Shao et al., 2015).

(B) A phase-contrast representative images of PTC and PDOs derived from surgically resected tumor samples of gastric and colorectal cancers at day 5. Scale bar: 100 µm. A similar number of tumor cells were initially used in both cases.

(C) Bar diagram showing the numbers of gastric or colorectal cancers-derived PTC successfully cultured from poorly, moderately, well or unspecified differentiated adenocarcinomas.

(D) Immunohistochemical staining of three types of markers, including the epithelial cell markers, CRC or GC biomarkers and stem cell/cell proliferation ones. Scale bar: 100 µm.

(E) Immunofluorescent co-staining of day 5 PTC for the epithelial markers CK8/CK18 and the fibroblast marker FAP. Nuclear counterstaining (blue, Hoechst).

(F) Quantitative analysis of the proportion of epithelial cells (CD326+), fibroblast cells (FAP+) or macrophage cells (CD45+ or CD 11b+) in five PTC samples by flow cytometry.

#### Figure S2

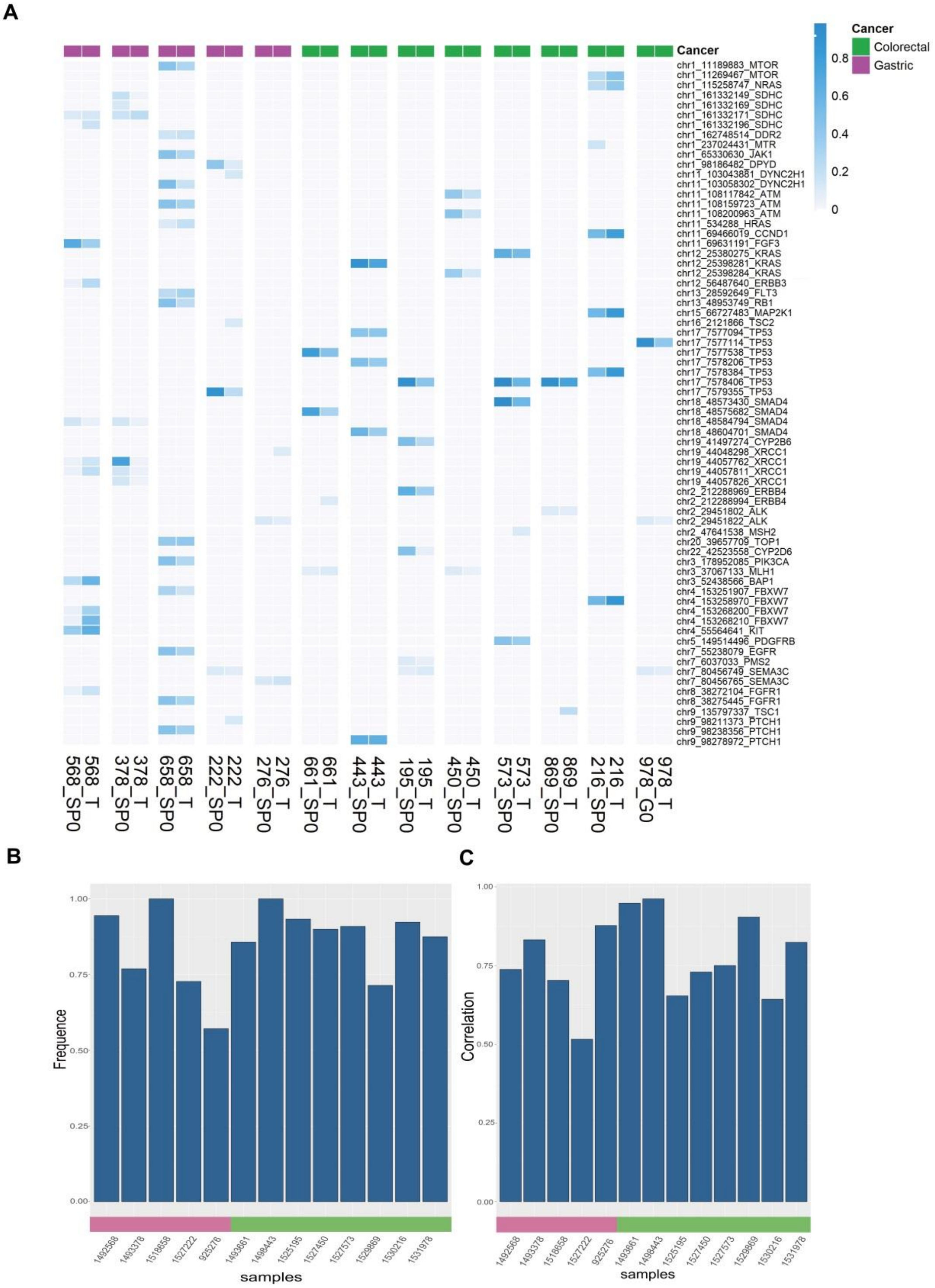


**Figure S2. Genomic Comparison of PTC with the Original Tumors, Related to Figure 2.**

Five gastric samples and eight colorectal samples were used to generate PTC. The last three digits of the sample names in B and C were used to name the corresponding samples in A. T: tumor samples; Sp: spheres of PTC.

(A) The VAFs of somatic in PTC and tumor biopsies. Synonymous mutations were not shown.

(B) The barplot of proportions of common somatic mutations in PTC and tumor biopsies.

(C) The barplot of the Pearson correlations of CNVs between PTC and tumor biopsy.

#### Figure S3

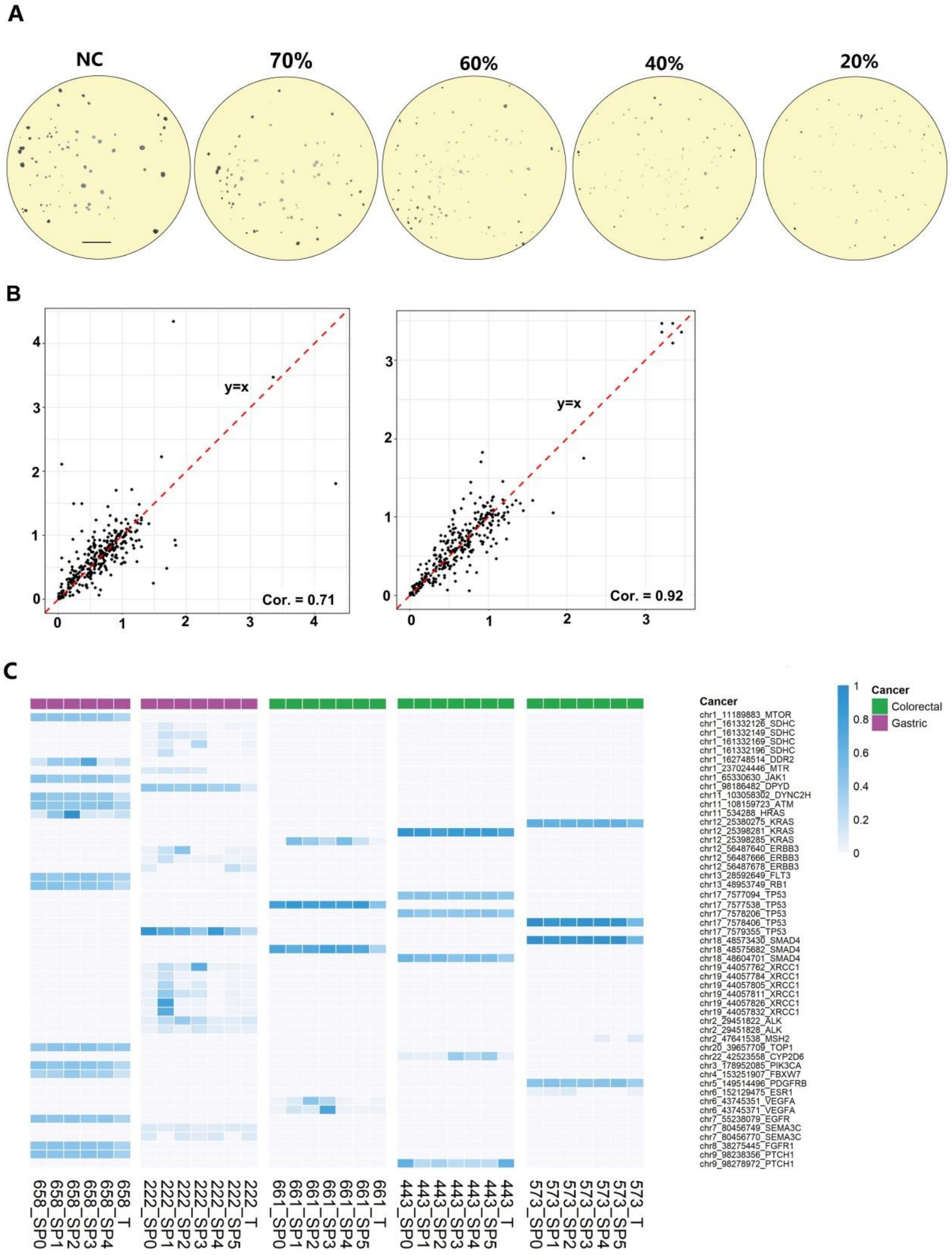
A
NC
70%
60%
40%
20%
B
y=x
Cor. = 0.71
y=x
Cor. = 0.92
C
Cancer
chr1_11189883_MTOR
chr1_161332126_SDHC
chr1_161332149_SDHC
chr1_161332169_SDHC
chr1_161332196_SDHC
chr1_162748514_DDR2
chr1_237024446_MTR
chr1_65330630_JAK1
chr1_98186482_DPYD
chr11_103058302_DYNC2H
chr11_108159723_ATM
chr11_534288_HRAS
chr12_25380275_KRAS
chr12_25398281_KRAS
chr12_25398285_KRAS
chr12_56487640_ERBB3
chr12_56487666_ERBB3
chr12_56487678_ERBB3
chr13_28592649_FLT3
chr13_48953749_RB1
chr17_7577094_TP53
chr17_7577538_TP53
chr17_7578206_TP53
chr17_7578406_TP53
chr17_7579355_TP53
chr18_48573430_SMAD4
chr18_48575682_SMAD4
chr18_48604701_SMAD4
chr19_44057762_XRCC1
chr19_44057784_XRCC1
chr19_44057805_XRCC1
chr19_44057811_XRCC1
chr19_44057826_XRCC1
chr19_44057832_XRCC1
chr2_29451822_ALK
chr2_29451828_ALK
chr2_47641538_MSH2
chr20_39657709_TOP1
chr22_42523558_CYP2D6
chr3_178952085_PIK3CA
chr4_153251907_FBXW7
chr5_149514496_PDGFRB
chr6_152129475_ESR1
chr6_43745351_VEGFA
chr6_43745371_VEGFA
chr7_55238079_EGFR
chr7_80456749_SEMA3C
chr7_80456770_SEMA3C
chr8_38275445_FGFR1
chr9_98238356_PTCH1
chr9_98278972_PTCH1
Cancer
Colorectal
Gastric
658_SP0
658_SP1
658_SP2
658_SP3
658_SP4
658_T
222_SP0
222_SP1
222_SP2
222_SP3
222_SP4
222_SP5
222_T
661_SP0
661_SP1
661_SP2
661_SP3
661_SP4
661_SP5
661_T
443_SP0
443_SP1
443_SP2
443_SP3
443_SP4
443_SP5
443_T
573_SP0
573_SP1
573_SP2
573_SP3
573_SP4
573_SP5
573_T

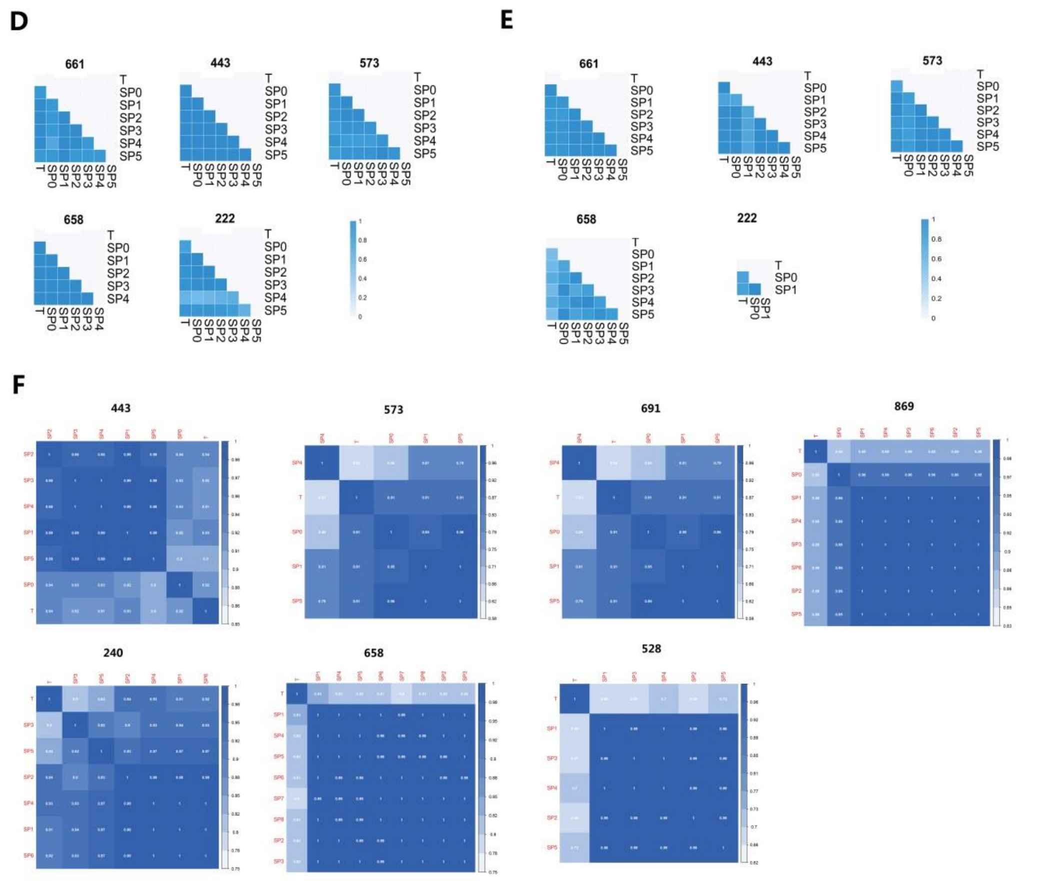


**Figure S3 Drug and Genomic Consistencies of PTC in Different Wells, Related to Figure 3.**

**(A-B) Drug Consistencies of PTC in Different Wells.** (A) An illustration of the five categories of drug efficacy. No effect: $p > 0.7$; weak effect: $0.7 > p \geq 0.6$; modest effect: $0.6 > p \geq 0.4$; significant effect: $p < 0.4$; and strong effect: $p < 0.2$. Only clusters with diameters greater than 40 micrometers were selected to calculate the areas. Scale bar: 1 mm. (B) Scatter diagrams showing the gastrointestinal cancer cases with the worst and best Pearson correlations. Cor. = 0.71 (left) and Cor. = 0.92 (right).

**(C-E) Somatic Mutations and CNV Consistencies of PTC in Different Wells.** Similar to Figure S2A, the last three digits of the sample names in Figure S2B and S2C were used to name the corresponding samples in S3. T: tumor samples; Sp: spheres or clusters of PTC. 0-5 means the PTC in different wells. (C) The VAFs of somatic mutations in replicates of PTC and tumor samples. Synonymous mutations were excluded. (D) The heatmaps of proportions of common somatic mutations among pairs of PTC and tumor samples. (E) The heatmaps of the Pearson correlation coefficients of CNVs among pairs of PTC and tumor samples.

**(F) Transcriptome Consistencies of PTC in Different Wells**. The heatmaps of the gene expression correlations among pairs of PTC and tumor biopsies.

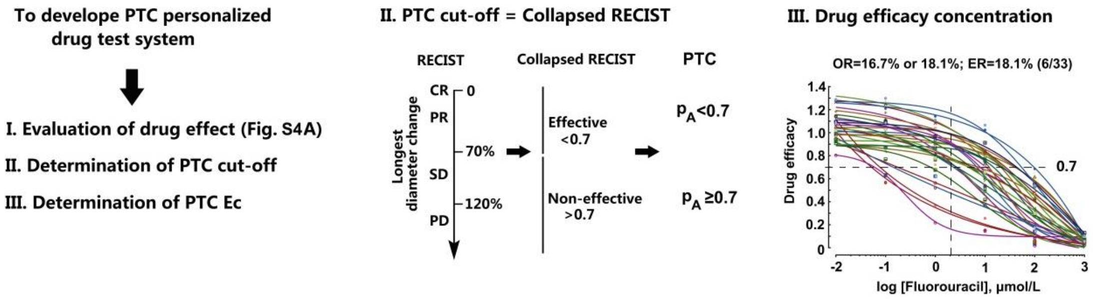


**Figure S4. The establishment of the PTC system for personalized drug testing**

Three key issues involved in the development of PTC system. The drug effect was evaluated by measuring the area change of clusters (Figure S4A). Cut-off and efficacy concentration of a drug in PTC drug testing are determined by its clinical efficacy. Cut-off is consistent with that of the collapsed RECIST, which is 0.7. Efficacy concentration (Ec) is determined by when the *in vitro* efficacy of the drug is consistent with the overall clinical response rate. For example, fluorouracil dose-response curves were created for PTCs generated from 33 GC and CRC patients using a 6-parameter logistic model; Ec was determined to be 2 μmol/L, since its efficacy rate was 18.1% (6/33) while the overall response rate of fluorouracil in CRC clinical trials was 16.7% or 18.1%.

**Materials and Methods**

**Human tumour samples**

All human tissue samples were obtained from Beijing Cancer Hospital. Before surgery at the center, all patients provided written informed consent to allow any excess tissue to be used for research studies. The study was approved by the ethical committee. Samples were snap frozen in OCT and stored at 0 ℃ until use. The pathologic status of the specimens was provided by the hospital.

**Patient population**

Enrollment criteria:

1. Adult patients between 18 and 75 years old, male or female;
2. Voluntary patient consent；
3. Study subjects are pathology confirmed, stage III/IV gastrointestinal cancer patients who underwent surgery to remove tumor tissue or biopsy, and can tolerate chemotherapy with measurable tumor lesion or malignant ascites.
4. ECOG status <3；
5. Patient has at least one measurable disease lesion (according to RECIST1.1).

Exclusion criteria:

1. Participated in any other clinical study within 6 months;
2. Women currently breast feeding or pregnant;
3. Severe liver or kidney function impairment (Live function: TBIL ＞1.5×ULN，ALT & AST＞2.5×ULN);Kidney function：Cr ＞1.5×ULN and creatinine clearance rate≥ 50 mL/min (according to the Cockcroft-Gault formula);
4. Patients with cognitive impairment, psychological disease, or poor compliance;
5. Allergic to known chemotherapy ingredients;

Other factors researchers deemed not suitable for study participation.

**Culture of PTCs from solid tumors**

Collected samples were conditioned in ice cold PBS with 10mM HEPES and 100 U/mL Penicillin-streptomycin (Thermo Fisher Scientific). Tissues were washed with PBS five times at least. Necrotic area and adipose tissue were removed as possible. Tissues were minced into small pieces and digested in 5 ml PBS/EDTA 1 mM containing collagenase I, II, and IV (Thermo Fisher Scientific) 200 U/mL each, at 37 ℃ for 1 hour. Tissues were minced into small pieces and digested in 5 ml PBS/EDTA 1 mM containing 2x TrypLe (Thermo Fisher Scientific) 200 U/mL each, at 37 ℃ for 1 hour. Pipetting every 15 minutes to facilitate cell release. 40 μm filters were used to collect dissociated cells. After 10 minutes' centrifugation (300 × g, 4 °C), cell pellets were resuspended in PTC growth medium, and seeded in low-attachment-surface dish at the concentration of $10^5$ cells/cm$^2$. Cells were cultured in incubator at 37 ℃, 5% $CO_2$. PTC growth medium was refreshed every 2-3 days, as necessary.

**Culture of PTCs from ascites**

Ascites samples were conditioned on ice. Samples were centrifuged at 300 × g and 4 °C for 5minutes. Cell pellets were resuspended in PBS. Resuspended cells were added over Lymphocyte Separation Medium (MPbio) in centrifuge tube. After 20 minutes' centrifugation (400 × g, 4 °C), middle cell layer was collected, resuspended in PTC growth medium, and seeded in low-attachment-surface dish at the concentration of $10^5$ cells/cm$^2$. Cells were cultured in incubator at 37 ℃, 5% $CO_2$. PTC growth medium was refreshed every 2-3 days, as necessary.

**Passage of PTCs**

The PTCs were digested using TrypLe (Thermo Fisher Scientific). Briefly, the PTCs were dissociated into single cells by pipetting every 2 minutes. After they were washed with DMEM (Thermo Fisher Scientific), the cells were centrifuged at 300 × g and 4 °C for 10 minutes. The pellets were resuspended in PTC growth medium and seeded at a concentration of $10^5$ cells/cm$^2$.

**Gastrointestinal (GI)-PTC growth medium**

GI-PTC were cultured in Advanced DMED containing 1mM HEPES (Thermo Fisher Scientific), 1x GlutaMAX (Thermo Fisher Scientific), 100 U/mL Penicillin-streptomycin (Thermo Fisher Scientific), 1x B27 (Thermo Fisher Scientific), 1x Non-essential amino acids (Thermo Fisher Scientific) and the following additives.

| Additive | Supplier | Cat. No. | Concentration |
|---|---|---|---|
| EGF | Peprotech | AF-100-15 | 40 ng/mL |
| FGF-basic | Peprotech | 100-AF-18B | 20 ng/mL |
| TGFβ | Invitrogen | PHG9214 | 0.5 ng/mL |
| R-spondin 1 | Novoprotein Scientific | CD83 | 500 ng/mL |
| Noggin | Novoprotein Scientific | C018 | 100 ng/mL |
| Nicotinamide | Sigma-Aldrich | N0636 | 10 mM |
| GX-83-01-03 | GeneX Health | GX-C-03 | 0.5 µM |
| GX-202190-03 | GeneX Health | GX-C-03 | 10 µM |
| GX-27632-07 | GeneX Health | GX-C-07 | 10 µM |
| N-acetyl-L-cysteine | Sigma-Aldrich | A9165 | 1 mM |
| HGF | PeproTech | 100-39 | 30 ng/ml |
| MSP | Invitrogen | PIRGMCSF20 | 5 ng/mL |
| Prostaglandin E2 | R&D system | 2296 | 10nM |
| Primocin | InvivoGen | ant-pm-1 | 100mg/ml |
| CHIR99021 | Stemgent | 04-0004 | 20 µM |
| Wnt-3a | R&D | 5036-WN | 250 ng/mL |
| Gastrin | NJpeptide | 39024-57-2 | 10 nM |

**PTC drug assay**

The PTC drug screens were conducted in 96-well cell culture plates following the procedure described previously (Pauli et al., 2017; Vlachogiannis et al., 2018). Briefly, PTC that were more than 40 µm in diameter were collected using 40-µm filters (BD Falcon), centrifuged at 300 × g and 4 °C for 10 minutes, washed with PBS, and resuspended with G/I-PTC growth medium. Then, 100 µL of a medium containing 30-50 PTC was seeded into a Teflon-modified chip (GeneX Health, GX-01).

Next, 50 μL of PTC growth medium containing the drug was added to the well. Images of each well were screened with the Nikon Ti-U microscope system. The plates were incubated at 37 ℃ and 5 % $CO_2$. After drug treatment, the plates were screened again with the Nikon Ti-U microscope system. Compounds were sourced from commercial vendors and stored as 10 mM aliquots.

**PTC/tissue histology**

PTC were collected by centrifugation at 300 g and 4 ℃ for 10 minutes, washed with cold PBS, and fixed in PBS containing 4% paraformaldehyde overnight. The pellets were paraffin-embedded, and 5-μm-thick paraffin sections were generated according to the conventional method. The PTC or tissue morphology was determined by H&E staining following a standard staining protocol. The H&E images were obtained using a Nikon Ti2-U microscope.

**PTC immunohistochemistry**

Immunohistochemistry was performed as described previously (Srivastava et al., 2014). Briefly, paraffin sections were dewaxed and rehydrated through a graded ethanol series. After heat-mediated antigen retrieval with Citrate Unmasking Solution (Cell Signaling Technology), the sections were blocked at room temperature for 1 hour. The primary antibodies were diluted in 3% BSA, and staining was performed overnight at 4 ℃ with gentle rocking. DAB (Cell Signaling Technology) was used to provide an acceptable staining intensity. The stained tissue was visualized using the Nikon Ti2-U microscope.

**PTC immunofluorescence**

Immunofluorescence was performed as described previously (Ahnfelt-Ronne et al., 2007). Briefly, the PTC were collected by centrifugation at 300 × g and 4 °C for 10 minutes, washed with cold PBS, and fixed in PBS containing 4% paraformaldehyde overnight. The fixed PTC were transferred to absolute MeOH for 1 h on ice and then incubated in Dent’s bleach (MeOH: DMSO: $H_2O_2$ = 4:1:1) for 2 h at room temperature. After equilibration to PBS through a series of descending MeOH concentrations in PBS (75%, 50%, and 25%), the samples were blocked using 3% BSA in PBS for 2 hours at room temperature. The primary antibodies were diluted in 3% BSA, and staining was performed overnight at 4 °C with gentle rocking. Alexa Fluor 488-labeled secondary antibodies were diluted in 3% BSA, and staining was performed at room temperature for 2 h. After 5 washes with PBS, the PTC were stained with Hoechst 33342 (10 μg/mL, Solarbio) for 15 min. The stained PTCs were visualized using a Zeiss LSM710 confocal microscope.

**Flow Cytometry**

PTC were harvested with 40μm filters, and digested into single cells using TrypLe (Thermo Fisher Scientific). After centrifugation, a single cell suspension was prepared in Cell Staining Buffer (Biolegend). Cells were centrifuged at 350×g for 5 minutes, and pellets were resuspended in Cell Staining Buffer at $10^6$ cells/ml. Human TruStain FcX™(Biolegend) was used to block Fc receptors. Conjugated fluorescent antibodies were added and cell suspensions were incubated on ice for 15 minutes in the dark. Cells were washed twice with Cell Staining Buffer and then analyzed with a Flow Cytometer (BD).

**Immunofluorescence (IF) and immunohistochemistry (IHC)**

The primary antibodies used in immunohistochemistry and immunofluorescence assays are listed in the table below:

| Antibodies | Supplier | Cat. No. | Application | Dilution |
|---|---|---|---|---|
| Galetin-3 | Cell Signaling | 87985 | IHC, IF | IHC 1:2000<br>IF 1:1000 |
| CK8/CK18 | Abcam | ab17139 | IHC, IF | IHC 1:50<br>IF 1:100 |
| S100A4 | Abcam | ab124805 | IHC, IF | IHC 1:400<br>IF 1:200 |
| GATA-3 | Cell Signaling | 5852 | IHC, IF | IHC 1:2000<br>IF 1:1000 |
| CDX2 | Abcam | ab76541 | IHC, IF | IHC 1:400<br>IF 1:200 |
| CD133 | Abcam | ab19898 | IHC, IF | IHC 1:200<br>IF 1:1000 |
| PDX1 | Abcam | ab47267 | IHC, IF | IHC 1:5000<br>IF 1:2000 |
| TTF-1 | Abcam | ab76013 | IHC, IF | IHC 1:250<br>IF 1:500 |
| Vimentin | Abcam | ab92547 | IHC, IF | IHC 1:300<br>IF 1:300 |
| MUC2 | Abcam | ab11197 | IHC, IF | IHC 1:600<br>IF 1:300 |
| SOX2 | Santa Cruz | sc-365823 | IHC, IF | IHC 1:500<br>IF 1:100 |
| CD11b | Abcam | ab133357 | IHC, | 1：4000 |
| Villin | Abcam | ab130751 | IHC | 1:100 |
| ALDH1A1 | Cell Signaling | 54135 | IHC | 1:200 |
| α-SMA | Thermo | 701457 | IHC | 1:500 |
| NapsinA | Abcam | 5640ab73021 | IHC | 1:100 |
| Ki-67 | Cell Signaling | 9129 | IF | 1:400 |
| ALDH1A1 | Cell Signaling | 36671 | IF | 1:400 |
| E-Cadherin | Cell Signaling | 3195 | IF | 1:500 |
| CD44 | Cell Signaling | 5640 | IF | 1:400 |
| SOX2 | Abcam | ab97959 | IF | 1:500 |
| CD326 | Biolegend | 324203 | Flow Cyto | 5μL/$10^6$ cells |
| CD45 | Biolegend | 368509 | Flow Cyto | 5μL/$10^6$ cells |
| CD11b | Biolegend | 301405 | Flow Cyto | 5μL/$10^6$ cells |
| FAP | R&D | FAB3715R | Flow Cyto | 1μL/$10^6$ cells |

**Cell Viability Assay**

For PTCs/Spheroids in a 96-well, media was half removed and replaced with 80 μL Celltiter-Glo cell viability assay (Promega). Cells were incubated at room temperature for 1 hour, and viability readings were obtained using Synergy plate reader (BioTek).

**EdU Assay**

Cell proliferation ability was measured by Click-it EdU imaging kits (Life, Alexa Flour 594). In brief, PTCs or Spheroids were incubated in growth medium with 20 μM EdU for 1 hour. After a 4% PFA fixation step and a 0.5% Triton X-100 permeabilization step, cells were incubated in Click-iT Plus reaction cocktail for 30 minutes. Nucleuses were marked by 5 μg/mL Hoechst 33342 for 4 hours. Images were captures with Nikon Ti-U microscope system.

**DNA and RNA extraction**

For samples preparation, the specimens were washed twice in phosphate buffered saline (PBS) prior to DNA or RNA isolation. DNA was extracted from tissues or cultured cells by proteinase K digestion in combination with the DNeasy Blood & Tissue Kit (Qiagen), following the protocol. RNA was isolated using Trizol method. Each sample was re-suspended in 500 μL Trizol (thermo Fisher Scientific), briefly vortexed and 100 μL Chloroform (sigma) was added. Phase separation was achieved by centrifuging the sample for 12,000 × g for 15 minutes at 4 °C, 200 μL of the aqueous phase was carefully transferred to a new tube. The RNA precipitation was carried out with 200μL Isopropanol (sigma) and the RNA pellet was washed with 1 mL of 70% ice cold ethanol.

**DNA and RNA library preparation and sequencing**

Target library preparation and sequencing were outsourced to GX-Health corporation (China). In brief, Whole genomic DNA libraries were carried out using NEBNext Ultra II DNA library Prep Kit for Illumina (New England Biolabs), the protocol consists of several enzymatic and purification steps. DNA target sequencing libraries were generated with the Cancer panel that targets 105 cancer-related genes. RNA libraries were prepared by NEBNext Ultra II RNA library Prep Kit for Illumina (New England Biolabs). Libraries were analyzed by Bioanalyzer 2100 and DNA 1000 or DNA high sensitivity chips, to quantify the library size and assess the level of adapter-dimer and primer-dimer contamination. Libraries were quantified using Illumina library qPCR quantification kits from KAPA Biosystems. Paired-end sequencing (2x150bp) was performed sequenced with Illumina HiSeq Xten.

**Genomic mutation and copy number analysis, mRNA sequencing analysis**

Mean depth of target sequencing samples was 967X (range from 211X to 1957X). Paired-end reads were mapped to the human reference genome (hg19) using SpeedSeq (Chiang et al., 2015) pipeline. Five mutation calling methods, VarDict (Lai et al., 2016), LoFreq (Andreas et al., 2012), SiNVICT (Kockan et al., 2016), Pindel (Ye et al., 2009), FreeBayes (Garrison and Marth, 2012) and GATK (Mckenna et al., 2010), were used to detect SNVs and Indels. SNVs and Indels supported by at least two methods were selected to improve the mutation detection sensitivity and accuracy. We only kept SNVs/Indels with a minimum coverage 50 and a minimum of 5 supporting reads. Low variant allele frequency (<0.1) SNVs/Indels were also filtered. To remove germline mutations, we filtered SNVs/indels in PTC or tumor biopsies whose VAFs in the corresponding normal data were greater than 1% or which had over 2 supporting reads in the normal data. Then, SNVs/Indels of each patient were genotyped for all PTC and tumor tissue samples by Platypus v0.8.1 to recover potential missed mutations in the calling step(Rimmer et al., 2014). Annovar (Wang et al., 2010) was used to annotate these SNVs. Please see supplementary **Table S4** for the all detected mutations, and **Table S5** for the overlaps of mutations between tumors and PTC.

Mean depth of whole genome sequencing was 3X (range 2X-5X). Paired-end reads were mapped to the human reference genome (hg19) using BWA (v0.7.12) (Li and Durbin, 2009a; Li and Durbin, 2009b). Possible PCR duplicates are removed using Picard tools and Samtools (Li et al., 2009) was used for sorting and indexing. BIC-seq2(Xi et al., 2016) was used to detect copy number variations (CNV). Regions with low mappability were filtered. This was achieved by removing regions with a low binNum and segment size ratio (<0.2) as reported by BIC-seq2. To compute CNV correlation of two samples, we first extracted all regions of both samples and split the region according to these breakpoints. Small sub-regions (<100kb) were removed and we calculated the Pearson correlation coefficient of the log2.copyratio reported by BIC-seq2 for the remaining sub-regions. Please see supplementary **Table S6** for BIC-seq2 segmentations, and **Table S7** for CNV correlations.

For mRNA sequencing analysis, paired-end reads were aligned to the reference genome(hg19) using Hisat2 (v2.1.0) (Kim et al., 2015). The FPKM value for each gene of each sample was estimated with StringTie (v1.3.3b) (Pertea et al., 2015) and the Pearson correlation were calculated according to these FPKM values.

**Of note, the sample names in the main manuscript are the last three digits of the sample names in supplementary Table S3-S6**.

**Statistical analysis**

A total of n = 3 independent experiments were conducted unless otherwise stated. n.s. indicates a nonsignificant difference. The P values were calculated using the unpaired two-tailed Student's t-test and reported as *P < 0.05, **P < 0.01, ***P < 0.001, and ****P < 0.0001.

#### Reference


Ahnfelt-Ronne, J., Jorgensen, M. C., Hald, J., Madsen, O. D., Serup, P., and Hecksher-Sorensen, J. (2007). An improved method for three-dimensional reconstruction of protein expression patterns in intact mouse and chicken embryos and organs. The journal of histochemistry and cytochemistry : official journal of the Histochemistry Society *55*, 925-930.

Andreas, W., Kim, A. P. P., Denis, B., Ting, Y. G. H., Hoe, O. S., Hua, W. C., Chuen, K. C., Rosemary, P., Lloyd, H. M., and Niranjan, N. (2012). LoFreq: a sequence-quality aware, ultra-sensitive variant caller for uncovering cell-population heterogeneity from high-throughput sequencing datasets. Nucleic Acids Research *40*, 11189.

Chiang, C., Layer, R. M., Faust, G. G., Lindberg, M. R., Rose, D. B., Garrison, E. P., Marth, G. T., Quinlan, A. R., and Hall, I. M. (2015). SpeedSeq: ultra-fast personal genome analysis and interpretation. Nature Methods *12*, 966.

Cunningham, D., Zalcberg, J. R., Rath, U., Oliver, I., van Cutsem, E., Svensson, C., Seitz, J. F., Harper, P., Kerr, D., and Perez-Manga, G. (1996). Final results of a randomised trial comparing 'Tomudex' (raltitrexed) with 5-fluorouracil plus leucovorin in advanced colorectal cancer. "Tomudex" Colorectal Cancer Study Group. Ann Oncol *7*, 961-965.

Eisenhauer, E. A., Therasse, P., Bogaerts, J., Schwartz, L. H., Sargent, D., Ford, R., Dancey, J., Arbuck, S., Gwyther, S., Mooney, M., *et al.* (2009). New response evaluation criteria in solid tumours: revised RECIST guideline (version 1.1). Eur J Cancer *45*, 228-247.

Garrison, E., and Marth, G. (2012). Haplotype-based variant detection from short-read sequencing. Quantitative Biology.

Kim, D., Langmead, B., and Salzberg, S. L. (2015). HISAT: a fast spliced aligner with low memory requirements. Nature Methods *12*, 357-360.

Kockan, C., Hach, F., Sarrafi, I., Bell, R. H., Mcconeghy, B., Beja, K., Haegert, A., Wyatt, A. W., Volik, S. V., and Chi, K. N. (2016). SiNVICT: Ultra-Sensitive Detection of Single Nucleotide Variants and Indels in Circulating Tumour DNA. Bioinformatics *33*, 26.

Lai, Z., Markovets, A., Ahdesmaki, M., Chapman, B., Hofmann, O., Mcewen, R., Johnson, J., Dougherty, B., Barrett, J. C., and Dry, J. R. (2016). VarDict: a novel and versatile variant caller for next-generation sequencing in cancer research. Nucleic Acids Research *44*, e108-e108.

Li, H., and Durbin, R. (2009a). Fast and accurate long-read alignment with Burrows–Wheeler transform.

Li, H., and Durbin, R. (2009b). Fast and accurate short read alignment with Burrows-Wheeler transform.

Li, H., Handsaker, B., Wysoker, A., Fennell, T., Ruan, J., Homer, N., Marth, G., Abecasis, G., and Durbin, R. (2009). The Sequence Alignment/Map (SAM) Format and SAMtools. Transplantation Proceedings *19*, 1653-1654.

Liu, Y., Wu, W., Hong, W., Sun, X., Wu, J., and Huang, Q. (2014). Raltitrexed-based chemotherapy for advanced colorectal cancer. Clin Res Hepatol Gastroenterol *38*, 219-225.

Mckenna, A., Hanna, M., Banks, E., Sivachenko, A., Cibulskis, K., Kernytsky, A., Garimella, K., Altshuler, D., Gabriel, S., and Daly, M. (2010). The Genome Analysis Toolkit: a MapReduce framework for analyzing next-generation DNA sequencing data. Genome Research *20*, 1297-1303.

Ohtsu, A., Shimada, Y., Shirao, K., Boku, N., Hyodo, I., Saito, H., Yamamichi, N., Miyata, Y., Ikeda, N., Yamamoto, S., *et al.* (2003). Randomized phase III trial of fluorouracil alone versus fluorouracil plus cisplatin versus uracil and tegafur plus mitomycin in patients with unresectable, advanced gastric cancer: The Japan Clinical Oncology Group Study (JCOG9205). J Clin Oncol *21*, 54-59.

Pauli, C., Hopkins, B. D., Prandi, D., Shaw, R., Fedrizzi, T., Sboner, A., Sailer, V., Augello, M., Puca, L., Rosati, R., *et al.* (2017). Personalized In Vitro and In Vivo Cancer Models to Guide Precision Medicine. Cancer Discov *7*, 462-477.

Pertea, M., Pertea, G. M., Antonescu, C. M., Chang, T. C., Mendell, J. T., and Salzberg, S. L. (2015). StringTie enables improved reconstruction of a transcriptome from RNA-seq reads. Nature Biotechnology *33*, 290-295.

Rimmer, A., Hang, P., Mathieson, I., Iqbal, Z., Twigg, S. R. F., Consortium, W., Wilkie, A. O. M., Mcvean, G., and Lunter, G. (2014). Integrating mapping-, assembly- and haplotype-based approaches for calling variants in clinical sequencing applications. Nature Genetics *46*, 912-918.

Shao, Z., Zhang, X., Pi, Y., Yin, L., Li, L., Chen, H., Zhou, C., and Ao, Y. (2015). Surface modification on polycaprolactone electrospun mesh and human decalcified bone scaffold with synovium-derived mesenchymal stem cells-affinity peptide for tissue engineering. J Biomed Mater Res A *103*, 318-329.

Shi, W. L., Sun, M. Y., Hu, X. Q., Ren, B., Cheng, J., Li, C. X., Duan, X. N., Fu, X., Zhang, J. Y., Chen, H. F., and Ao, Y. F. (2017). Structurally and Functionally Optimized Silk-Fibroin-Gelatin Scaffold Using 3D Printing to Repair Cartilage Injury In Vitro and In Vivo. Adv Mater *29*.

Srivastava, S., Liew, M. S., McKeon, F., Xian, W., Yeoh, K. G., Ho, K. Y., and Teh, M. (2014). Immunohistochemical analysis of metaplastic non-goblet columnar lined oesophagus shows phenotypic similarities to Barrett's oesophagus: a study in an Asian population. Digestive and liver disease : official journal of the Italian Society of Gastroenterology and the Italian Association for the Study of the Liver *46*, 170-175.

Vlachogiannis, G., Hedayat, S., Vatsiou, A., Jamin, Y., Fernández-Mateos, J., Khan, K., Lampis, A., Eason, K., Huntingford, I., Burke, R., *et al.* (2018). Patient-derived organoids model treatment response of metastatic gastrointestinal cancers. Science *359*, 920-926.

Wang, K., Li, M., and Hakonarson, H. (2010). ANNOVAR: functional annotation of genetic variants from high-throughput sequencing data. Nucleic Acids Research *38*, e164.

Xi, R., Lee, S., Xia, Y., Kim, T. M., and Park, P. J. (2016). Copy number analysis of whole-genome data using BIC-seq2 and its application to detection of cancer susceptibility variants. Nucleic Acids Research *44*, 6274.

Ye, K., Schulz, M. H., Long, Q., Apweiler, R., and Ning, Z. (2009). Pindel: a pattern growth approach to detect break points of large deletions and medium sized insertions from paired-end short reads. Bioinformatics *25*, 2865.
